# PARP7 mono-ADP-ribosylates the Agonist Conformation of the Androgen Receptor in the Nucleus

**DOI:** 10.1101/2021.06.21.449333

**Authors:** Teddy Kamata, Chun-Song Yang, Bryce M. Paschal

## Abstract

We recently described a signal transduction pathway that contributes to AR regulation based on sitespecific ADP-ribosylation by PARP7, a mono-ADP-ribosyltransferase implicated in several human cancers. ADP-ribosylated AR is specifically recognized by PARP9/DTX3L, a heterodimeric complex that contains an ADP-ribose reader (PARP9) and a ubiquitin E3 ligase (DTX3L). Here, we have characterized the cellular and biochemical requirements for AR ADP-ribosylation by PARP7. We found that the reaction requires nuclear localization of PARP7 and an agonist-induced conformation of AR. PARP7 contains a Cys_3_His_1_-type zinc finger (ZF), which we found is critical for AR ADP-ribosylation. The Parp7 ZF is required for efficient nuclear localization by the nuclear localization signal (NLS) encoded in PARP7, but rescue experiments indicate the ZF makes a contribution to AR ADP-ribosylation that is transport-independent. ZF structure appears to be dispensable for PARP7 catalytic activity and for PARP7 binding to AR. Androgen induction of the *MYBPC1* gene is regulated by AR and PARP7, and we determined that the ZF is required for the PARP7 transcriptional effect on *MYBPC1.* Our data indicate the PARP7 ZF plays an important role in modulating the subcellular localization of PARP7 and its capacity to ADP-ribosylate and promote AR-dependent transcription.

## Introduction

Over 1.2 million men worldwide were diagnosed in 2018 with prostate cancer (PCa), making it the second most frequently diagnosed cancer in men which, in addition, is associated with significant healthcare costs [1]. The primary driver of PCa is the androgen receptor (AR) signaling pathway. AR is a member of the nuclear receptor superfamily of ligand-activated transcription factors that binds androgen and adopts an active, agonist conformation [2,3]. The mainstay of PCa treatment is androgen deprivation therapy (ADT), the goal of which is to inhibit AR signaling by blocking androgen synthesis and/or preventing androgen binding to AR with anti-androgens that induce an inactive, antagonist conformation [4]. ADT generally achieves remission in PCa patients; however, disease relapse can occur as a result of reactivation of AR signaling pathway [5,6]. The failure of current therapies to manage PCa emphasizes the need for a better understanding of the regulatory mechanisms that control AR activity.

Recently, our lab described a novel signal transduction pathway that modulates AR activity through ADP-ribosylation [7]. The pathway is dependent on the mono-ADP-ribosyltransferase PARP7. The other names for this enzyme include 2.3.7.8-tetrachlorodibenzo-*p*-dioxin-inducible poly-ADP-ribose polymerase (TIPARP) and ADP-ribosyltransferase diphtheria toxin-like 14 (ARTD14). *PARP7* is a direct AR target gene and is induced by androgen in PCa cells [7,8]. Androgen treatment also promotes PARP7 protein stability through a mechanism that is AR-dependent but separable from *PARP7* gene induction [9]. These two mechanisms drive the expression of PARP7 and direct ADP-ribosylation of AR on multiple cysteine residues, which create binding sites for PARP9 (mono-ADP-ribosyltransferase)/DTX3L (E3 ubiquitin ligase) heterodimer. Recruitment of the PARP9/DTX3L complex to ADP-ribosylated AR is mediated through specific recognition of the ADP-ribose moiety by the two macrodomains within PARP9. PARP9/DTX3L complex assembly on AR appears to regulate the level of transcription output from a subset of genes in PCa cells. Thus, the AR-PARP7 signal transduction pathway provides an additional level of regulation for AR-dependent transcription.

Here, we characterize PARP7-mediated AR ADP-ribosylation and show the reaction is highly dependent on AR adopting an agonist conformation. We found that PARP7 localization to the nucleus is critical for AR ADP-ribosylation of AR, consistent with the reaction occurring within the nucleus. Our structure-function analysis showed that the Cys_3_His_1_-type zinc finger (ZF) domain in PARP7 modulates PARP7 nuclear import, but is also required for PARP7 ADP-ribosylation of AR.

## Results

### An agonist conformation of AR is required for ADP-ribosylation by PARP7

In previous work we showed that androgen regulates AR ADP-ribosylation through two, separable mechanisms that are both expression-based: androgen induction of *PARP7* transcription [7] and androgen induction of PARP7 protein stability [9]. Both mechanisms are critical for determining the cellular concentration of PARP7. Given that androgen induces conformational changes in AR, in the current study, we queried whether androgen binding might also promote ADP-ribosylation by modulating an AR structure in a manner that enhances its suitability as a substrate for PARP7. The precedent for androgen regulation of AR as a substrate derives from studies showing androgen induces multi-site phosphorylation [10]. To eliminate the aforementioned androgen effects on PARP7 expression that might complicate interpretation, ectopic PARP7 was expressed from a CMV promoter and assays were performed in HEK293T cells. We also employed two AR mutants that contain amino acid substitutions in the ligand binding domain (LBD). The L860F substitution (Figure 1A) was identified in a patient with complete androgen insensitivity syndrome and reduces androgen binding to ~14% of WT [11]. The T878A mutation commonly found as a resistance mechanism in PCa patients undergoing ADT broadens the ligand specificity of the LBD and enables the androgen antagonist hydroxyflutamide (HO-Flutamide) to induce an agonist conformation in AR (Figure 1A) [12–14].

**Figure 1.**
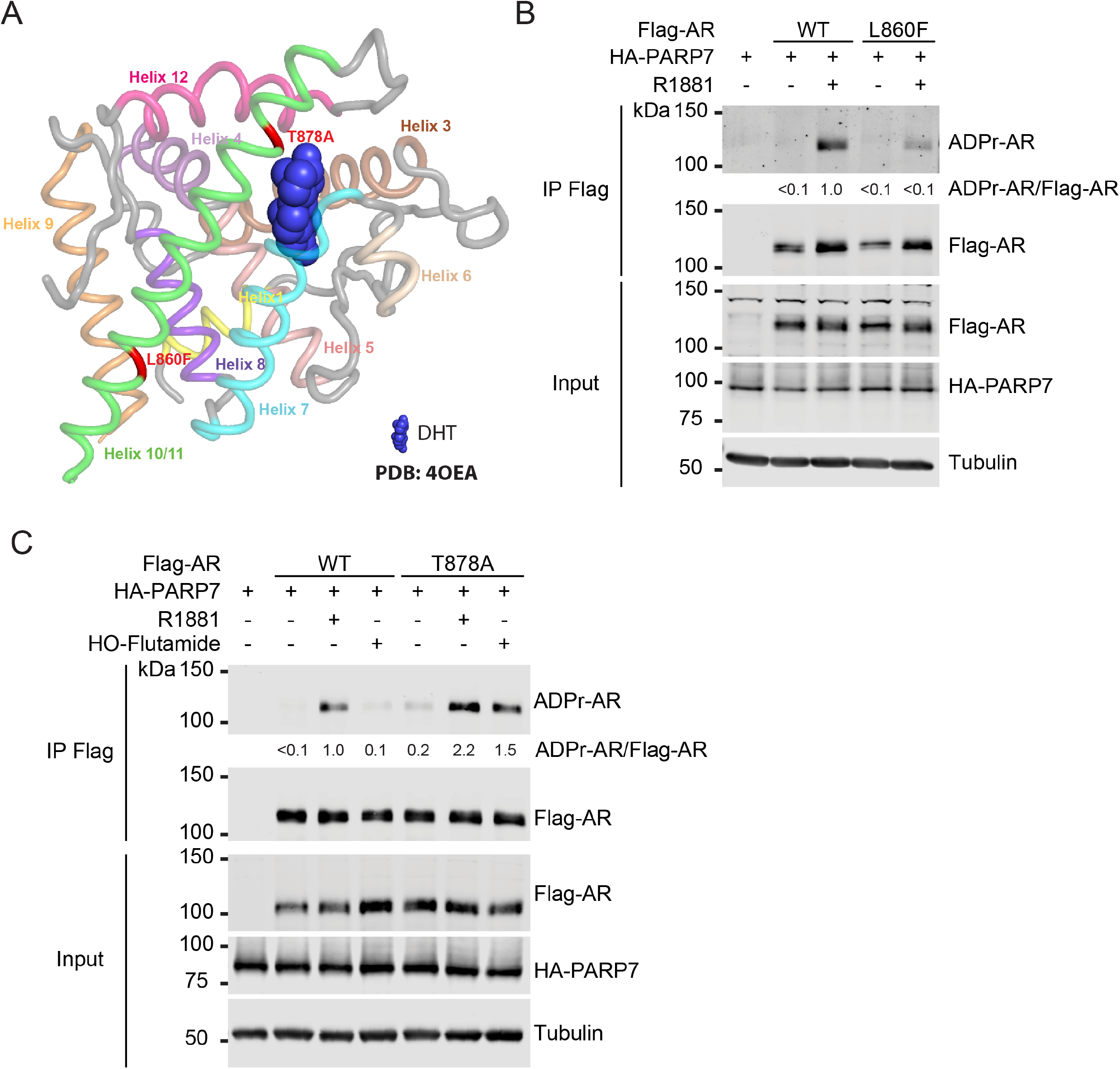
An agonist conformation of AR is required for ADP-ribosylation. (**A**) Structure of AR LBD bound to the agonist DHT (blue) (PDB: 4OEA). T877A and L860F substitutions are indicated in red. (**B**) HEK293T cells were transfected with HA-PARP7 and Flag-AR (WT or L860F). After 24 h, transfected cells were treated with R1881 for 8 h. Flag-AR was immunoprecipitated from treated cells and analyzed for ADP-ribosylation. ADP-ribosylated AR (ADPr-AR) was quantified as a ratio of ADPr-AR to AR. Ratios are displayed relative to the WT + R1881 lane (set to 1.0). (**C**) HEK293T cells were transfected with HA-PARP7 and Flag-AR (WT or T878A). After 24 h, cells were treated with R1881 or hydroxyflutamide (HO-Flutamide) for 8 h. Flag-AR was immunoprecipitated from treated cells and analyzed for AR ADP-ribosylation. ADPr-AR was quantified similar to B).

We transiently transfected HEK293T cells with HA-PARP7 alone, and with Flag-AR WT and Flag-AR L860F (Figure 1B). After treatment with vehicle or R1881 (synthetic androgen), AR was then immunoprecipitated (IP’d) and probed for ADP-ribosylation using fluorescently labeled AF1521 [9]. We found that co-expression of HA-PARP7 did not result in significant AR-ADP-ribosylation unless androgen was added to the cells (Figure 1B, lanes 2 and 3). Thus, androgen-induced changes in AR are required for ADP-ribosylation by PARP7. By contrast, androgen treatment of cells expressing HA-PARP7 and Flag-AR L860F showed only a low level of AR ADP-ribosylation (Figure 1B, lane 5). These results show that an androgen-induced conformation of AR is required for ADP-ribosylation. To directly compare PARP7 ADP-ribosylation of agonist and antagonist conformations of AR, we transfected cells with HA-PARP7 and Flag-AR WT, and treated cells with either R1881 (agonist) or the antiandrogen HO-Flutamide (antagonist) (Figure 1C). R1881 treatment induced a high level of AR ADP-ribosylation, while HO-Flutamide treatment showed very little AR ADP-ribosylation (Figure 1C, compare lanes 3 and 4). Using Flag-AR T878A, we found that treatment with HO-Flutamide resulted in AR ADP-ribosylation to a level comparable to WT AR treated with R1881 (Figure 1C, lanes 3 and 7). From these data, we conclude that efficient ADP-ribosylation of AR by PARP7 requires an androgen-induced, agonist conformation of AR. The HO-Flutamide induction of ADP-ribosylation of the T878A mutant corroborates this view since the drug selectively induce the agonist conformation in this mutant.

### PARP7 nuclear localization is required for ADP-ribosylation of AR

The subcellular distribution of PARP7 is context specific. PARP7 localizes to the cytoplasm in ovarian cancer cells [15], and its function in anti-viral response occurs in the cytoplasm [16,17]. By contrast, PARP7 is primarily nuclear in several settings including PCa cells (Bindesbøll et al., 2016; Gomez et al., 2018; Kamata et al., 2021; MacPherson et al., 2013; Roper et al., 2014). Because nuclear receptors such as AR shuttle between the nucleus and cytoplasm [18], PARP7 could encounter and ADP-ribosylate AR in the cytoplasm or the nucleus. We tested whether nuclear localization of PARP7 is required for AR ADP-ribosylation by using a series of nuclear transport signal mutations and fusions to regulate the steady state localization of PARP7 (Figure 2A). These included mutation of the endogenous PARP7 NLS (“AAA”), fusion with the SV40 NLS (“NLS”), and fusion of the c-Abl nuclear export signal (“NES”) in various combinations. Tet-inducible cell lines expressing the PARP7 nuclear transport signal mutants and fusions were generated and analyzed by immunofluorescence (IF) microscopy. WT PARP7 is predominantly nuclear in the absence of androgen, and the nuclear (N):cytoplasmic (C) ratio is further increased by treating cells with R1881 (Figure 2B, C). Nuclear localization of PARP7 is dependent on its NLS since the AAA mutant adopts a cytoplasmic distribution (Figure 2B, C). The nuclear import defect of the PARP7 AAA mutant is rescued by the SV40 NLS (Figure 2B,C). Finally, PARP7 can be forced into the cytoplasm with the c-Abl NES, and the effect is even more pronounced when combined with the AAA mutant (Figure 2B, C). We analyzed the effect of PARP7 localization on AR ADP-ribosylation by probing cell extracts for AR ADP-ribosylation and normalizing to the level of AR expression (Figure 2D). In this experiment, androgen induction of endogenous PARP7 results in a modest level of AR ADP-ribosylation which is increased by Tet induction of WT PARP7 (Figure 2D, compare lanes 2 and 4). While small effects of the transport signal fusions and mutations were detectable, the most striking effect on AR ADP-ribosylation was obtained with the NES-AAA PARP7 mutant. The NES-AAA PARP7 mutant, which is largely restricted to the cytoplasm, did not increase AR ADP-ribosylation above the level supported by endogenous PARP7 (Figure 2D, compare lanes 2 and 14). This result is consistent with PARP7 requiring nuclear localization in order to ADP-ribosylate AR. As expected, double label IF microscopy for PARP7 and AR shows that WT PARP7 and AR co-localize to the nucleus, but in cells expressing PARP7 NES-AAA and AR, the distribution is mutually exclusive in many cells (Figure 2E).

**Figure 2.**
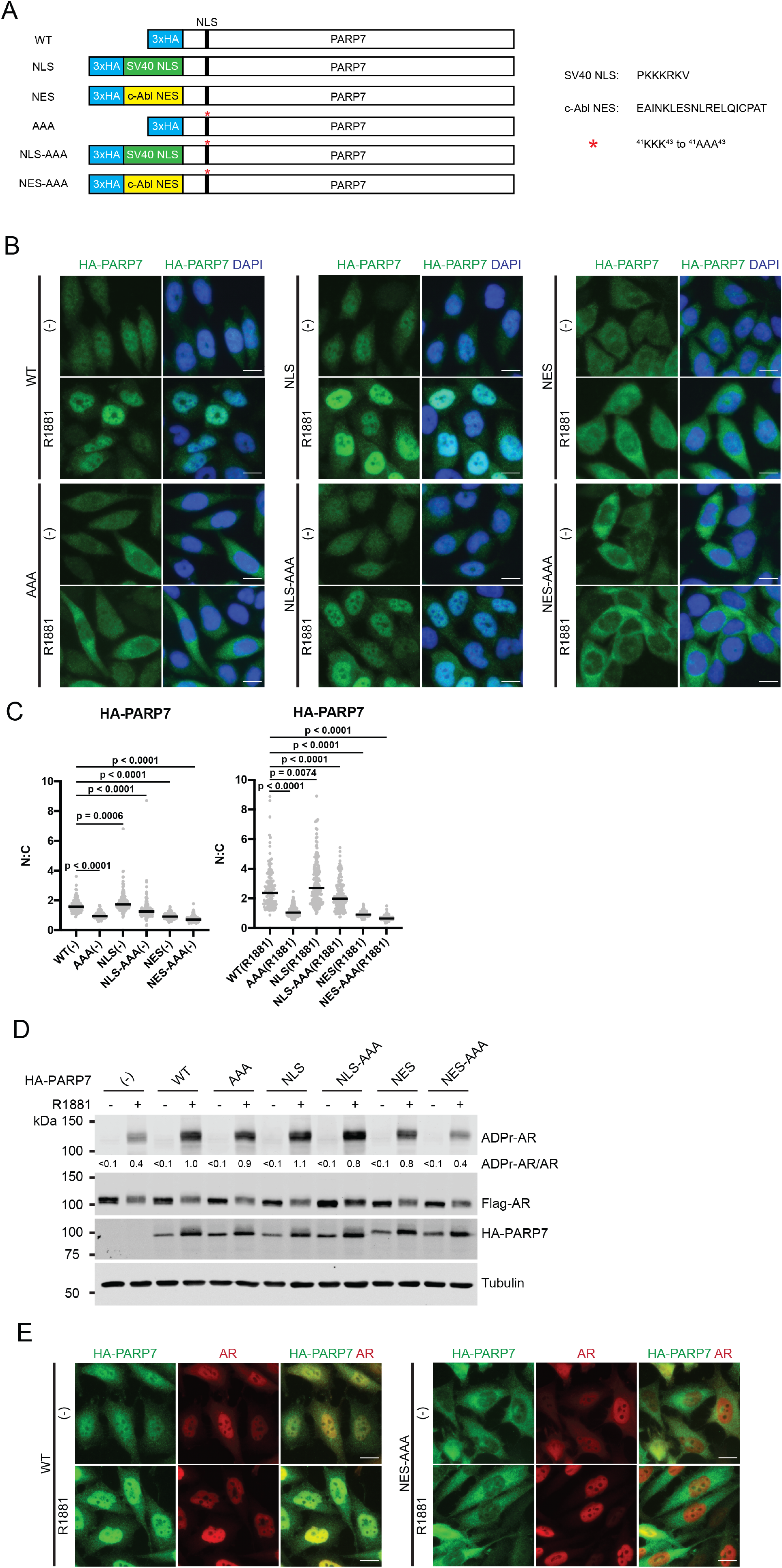
PARP7 nuclear localization is required for ADP-ribosylation of AR. (**A**) Schematic of PARP7 constructs used in this figure. As indicated, triple alanine subsitutions were made within the native NLS for PARP7 to generate the AAA PARP7 mutant. In addition, SV40 NLS or c-Abl NES were fused with WT or AAA PARP7. (**B**) HA-PARP7 WT or mutant were induced in PC3-Flag-AR cells by doxycycline (2 μg/mL, 24 h) and treated with R1881 for 6 h. Treated cells were processed for immunofluorescence microscopy. Scale bar = 5 μm. (**C**) Quantification of B). Distribution of HA-PARP7 was quantified as a ratio of nuclear (N) and cytoplasmic (C) signals for vehicle (left) and R1881-treated (right) groups. At least 100 cells were quantified for each condition, and median N:C ratio (black line) is indicated on the plots. One-way ANOVA with Tukey’s multiple comparison test was conducted to determine statistical significance. (**D**) PC3-Flag-AR cells expressing HA-PARP7 WT or mutant were induced with doxycycline and treated with R1881 as in B) and analyzed for AR ADP-ribosylation by immunoblotting. ADPr-AR was quantified as a ratio of ADPr-AR to AR. Ratios are displayed relative to the WT + R1881 lane (set to 1.0). (**E**) PC3-Flag-AR cells expressing WT or NES-AAA PARP7 were induced with doxycycline and treated with R1881 as in B). Treated cells were costained for HA-PARP7 and FlagAR for immunofluroescence microscopy. Scale bar = 5 μm.

### Zinc finger and catalytic domain of PARP7 are required for regulation of AR-dependent gene transcription

PARP7 encodes a single Cys_3_His_1_-type ZF that other groups have shown is required for the transcription regulatory effect of PARP7 on liver X receptors [19] and aryl hydrocarbon receptor [20]. Although the exact function of the PARP7 ZF in these studies was not defined, the data raises the interesting question of whether the ZF is generally required for PARP7 to affect transcription factors. The ZF (amino acids 237-264) is encoded after the N-terminal domain, which is predicted to be mostly unstructured (Figure 3A). We engineered single cysteine-to-alanine substitutions to test if the PARP7 ZF is important for AR ADP-ribosylation and the PARP7 effect on AR-dependent transcription, which includes assembly of the AR-PARP9/DTX3L complex [7]. As controls, we also generated loss-of-function point mutations in the PARP7 catalytic domain, which additionally allowed us to test for non-catalytic effects of PARP7 on the pathway. Using PC3-Flag-AR cells with Tet-inducible WT and mutant forms of PARP7, we assayed expression of the *MYBPC1* gene by RT-qPCR since it shows a strong dependence on PARP7 for efficient induction by androgen and AR [7]. In this assay, the level of *MYBPC1* expression in the absence of Dox but the presence of R1881 includes the contribution by endogenous PARP7 (Figure 3B, No Dox, black bar). We found that mutation of the ZF (C243A, C251A) eliminated the PARP7 enhancement of androgen-induced *MYBPC1* transcription (Figure 3B). Similarly, amino acid substitutions that inactivate the catalytic function of PARP7 (H332A, H532A) rendered PARP7 inactive for androgen-induced *MYBPC1* transcription (Figure 3B). Using bead-immobilized AF1521 to pull-down ADP-ribosylated proteins from cell extracts, we found that the ZF mutations in PARP7 reduce AR ADP-ribosylation, but both mutants (C243A, C251A) retain catalytic activity as indicated by PARP7 auto-modification (Figure 3C). The catalytic domain mutations in PARP7 eliminated AR ADP-ribosylation by ectopic PARP7, and these mutants showed only a low level of AF1521 binding (Figure 3C). We then examined the effects of the PARP7 mutations on AR complex formation with PARP9/DTX3L, a reaction that reflects PARP7 ADP-ribosylation of AR and PARP9/DTX3L binding via the macrodomain “reader” function of PARP9 [7]. Compared to WT PARP7, cell lines expressing the ZF and catalytic domain mutants of PARP7 showed only a low level of AR ADP-ribosylation and AR-PARP/DTX3L formation, and this was ascribable to endogenous PARP7 induction by R1881 (Figure 3D, lanes 2, 5-8). Taken together, the data show that the ZF and the catalytic domains of PARP7 are important for AR ADP-ribosylation, AR-PARP9/DTX3L complex formation, and enhancement of AR-dependent transcription of the *MYBPC1* gene.

**Figure 3.**
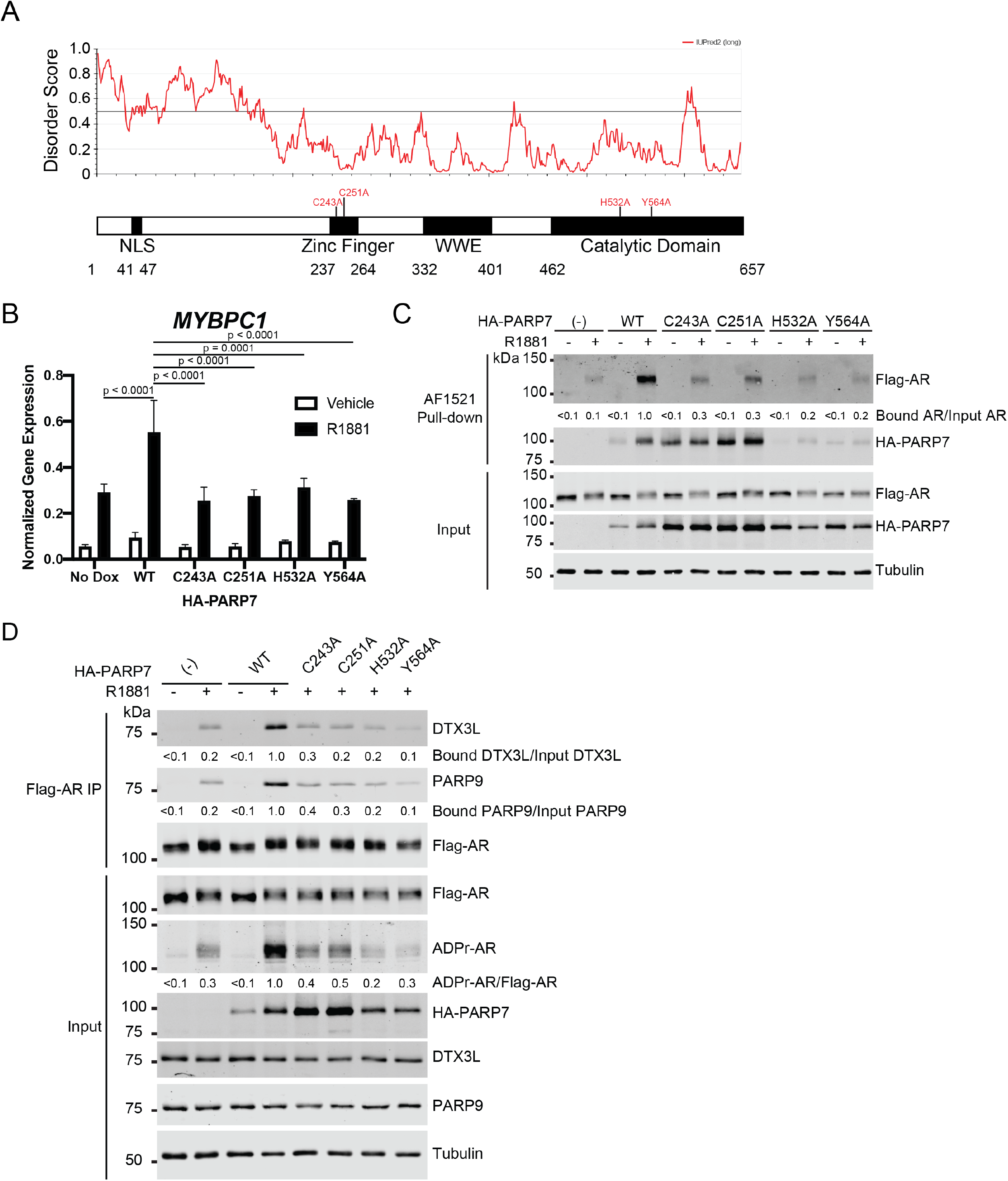
Both the zinc finger and catalytic domain within PARP7 are required for regulation of AR-dependent gene transcription through AR ADP-ribosylation and subsequent assembly of the PARP9/DTX3L complex. (**A**) Domain architecture of PARP7 with amino acid substitutions used throughout this study as well as plot of disorder regions within PARP7 predicted using IUPred2A [28]. NLS = nuclear localization signal, WWE = tryptophan-tryptophan-glutamate domain. (**B**) PARP7 WT or mutant (C243A, C251A, H532A, and Y564A) were induced in PC3-Flag-AR cells by doxycycline (2 μg/mL) for 24 hours. An untreated sample served as an uninduced control (No Dox). Cells were treated for 9 hours with androgen (2 nM R1881) before RNA was extracted and analyzed for gene expression of *MYBPC1.* Gene expression was normalized to the housekeeping gene *GUS.* Plots represent mean ± SD from three biological replicates. One-way ANOVA with Tukey’s multiple comparison test was conducted to determine statistical significance. (**C**) PC3-Flag-AR cells expressing HA-PARP7 WT or mutant were induced with doxycycline similar to B). Cells were subsequently treated with R1881 for 6 h, and AR and HA-PARP7 ADP-ribosylation was analyzed by AF1521 pull-down. Bound AR was quantified as a ratio of bound AR to input AR. Ratios are displayed relative to the WT + R1881 lane (set to 1.0). (**D**) PC3-Flag-AR cells expressing PARP7 WT or mutant were treated as in C). Flag-AR was immunoprecipitated from treated cells and analyzed for DTX3L and PARP9 complex assembly. Bound DTX3L and PARP9 was quantified similar to C). ADPr-AR was quantified as a ratio of ADPr-AR to input AR. Ratios are displayed relative to the WT + R1881 lane (set to 1.0).

### Zinc finger domain of PARP7 is neither sufficient nor required for interaction with AR

ZF function is usually associated with nucleic acid binding, but there are ZFs capable of mediating proteinprotein interactions [21]. Our finding that PARP7 ZF substitutions C243A and C251A reduced AR ADP-ribosylation led us to investigate if the ZF plays a role in AR binding. We addressed this question by co-expression of PARP7 and AR in HEK293T cells and AR IP, and immunoblotting. We found that the levels of PARP7 ZF mutant C243A and catalytic domain mutant H532A binding to AR was similar to that of PARP7 WT (Figure 4B). AR binding assays performed with deletion mutants of PARP7 (Figure 4A) showed that the catalytic domain of PARP7 is sufficient for the interaction (Figure 4C). PARP7 lacking a catalytic domain also was sufficient to bind AR, and this interaction was not reduced in the ZF C243A mutant (Figure 4C). These data are consistent with a model that PARP7 uses multiple domains to contact AR, and this is independent of ZF structure. We tested whether the PARP7 ZF is sufficient for binding AR using recombinant GST-ZF immobilized on beads and cell extracts that contain AR. We found no evidence of AR binding to the PARP7 ZF, or to the PARP7 ZF mutant C243A (Figure 4D). From these data we conclude the PARP7 ZF plays an important role in AR ADP-ribosylation that can be separated from protein binding and PARP7 catalytic activity.

**Figure 4.**
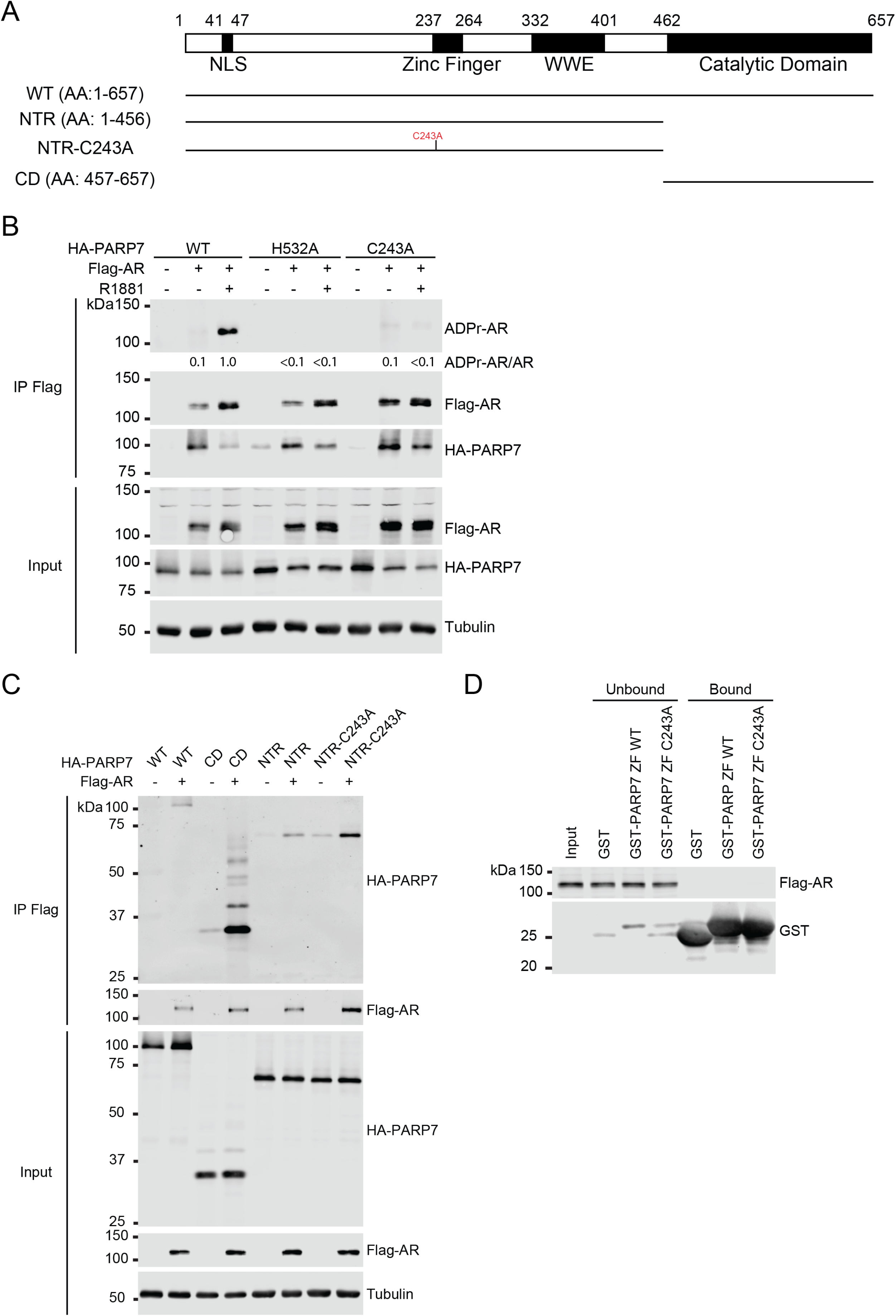
PARP7 zinc finger domain is neither sufficient nor required for interaction with AR. (**A**) A schematic of PARP7 deletion mutants used for co-immunopreciptiation studies in C). NTR = N-terminal region, CD = catalytic domain. (**B**) HEK293T cells were transfected with Flag-AR along with HA-PARP7 WT, H532A, or C243A. After 24 h, cells were treated with R1881 for 8 h. Flag-AR was immunoprecipitated from treated cells and analyzed for AR ADP-ribosylation and interaction with HA-PARP7. ADP-ribosylated AR (ADPr-AR) was quantified as a ratio of ADPr-AR to AR. Ratios are displayed relative to the WT + R1881 lane (set to 1.0). (**C**) HEK293T cells were transfected with the indicated HA-PARP7 construct either alone or with Flag-AR. After 24 h, Flag-AR was immunoprecipitated from transfected cells and analyzed for HA-PARP7 interaction. (**D**) Extract from PC3-Flag-AR cells treated with R1881 for 3 h was incubated with GST-fused PARP7 zinc finger domain (WT and C243A) immobilized on glutathione beads before analysis by SDS-PAGE and immunoblotting.

### Rescue of the nuclear localization defect for PARP7 zinc finger mutant does not restore AR ADP-ribosylation

To explore other mechanisms through which the PARP7 ZF contributes to AR ADP-ribosylation, we considered the observation by the Matthews group that the PARP7 ZF contributes to PARP7 nuclear localization [20,22]. Given that PARP7 requires nuclear localization to modify AR, mutations that reduce import would be predicted to reduce PARP7 ADP-ribosylation of nuclear substrates. We first examined the localization of WT and various PARP7 mutants in PC3-Flag-AR cells by IF microscopy. The preferential nuclear localization observed for WT PARP7 was lost when the C243A and C251A substitutions were introduced into the ZF (Figure 5A, B). Both the catalytic domain PARP7 mutants (H532A and Y564A) localized to the nucleus (Figure 5A,B), consistent with previous observations and emphasizing that PARP7 catalytic function is not a prerequisite for import [20,22,23]. With the goal of testing whether reduced AR ADP-ribosylation can be explained by defective import, we designed a rescue experiment that involved appending the SV40 NLS to the PARP7 C243A mutant.

We transfected WT, C243A, and NLS-C243A PARP7 (C243A PARP7 mutant fused to the SV40 NLS) into HEK293T cells, treated with androgen, and plotted PARP7 distribution as an N:C ratio signal from IF microscopy images. Our analysis showed that the C243A PARP7 mutant had a lower N:C ratio than that of WT PARP7, and that the NLS-C243A PARP7 construct had an N:C value that was not significantly different from WT PARP7 indicating rescue of the nuclear localization defect (Figure 5C,D). We then used these constructs to test whether rescue of nuclear localization restored AR ADP-ribosylation. Compared to WT PARP7, the PARP7 C243A mutant and the PARP7 NLS-C243A rescue construct showed only background levels of AR ADP-ribosylation (Figure 5E). Thus, although the ZF is important for efficient nuclear localization of PARP7, it appears to have a transport-independent function that is critical for AR ADP-ribosylation within the nucleus (Figure 5G).

Parenthetically, we noted about half of the cells transfected with WT PARP7 formed nuclear foci, a phenomena that has been observed by other groups (Figure 5C,F) [20,23]. PARP7 foci, which we observed in transfected HEK293T cells but not PCa cells, were not detected with either the PARP7 C243A or PARP7 NLS-C243A mutant, strongly indicating that PARP7 foci formation requires an intact ZF (Figure 5C,F).

## Discussion

In this study, we have expanded our understanding of PARP7/AR signaling by exploring the key characteristics within AR and PARP7 that are necessary for ADP-ribosylation of AR to occur. Our data underscore the point that agonist conformation of AR is required for ADP-ribosylation. The dependence on ligand binding for AR ADP-ribosylation is supported by the fact that the L860F mutation which disrupts androgen binding to AR [11] causes a substantial reduction in AR ADP-ribosylation. In HEK293T cells co-transfected with PARP7 and WT AR, we observed that androgen (agonist), but not HO-Flutamide (antagonist), treatment led to ADP-ribosylation of AR. The result can be explained by the fact that compared to agonist binding, AR adopts a distinct conformation when bound to antagonists such as bicalutamide and HO-Flutamide [24–26]. However, when the T878A mutation was introduced into the ligand binding domain of AR, ADP-ribosylation of AR was restored in the presence of HO-Flutamide. This is because the T878A AR mutant can adopt an agonist conformation even when the antagonist is bound [12]. From these results, we conclude that the agonist conformation for AR is a key requirement for ADP-ribosylation by PARP7.

Another key finding from our study is the requirement of an intact PARP7 ZF domain for ADP-ribosylation of AR and concomitant assembly of the PARP9/DTX3L complex and regulation of AR-dependent transcription. Neither the loss of catalytic activity nor loss of substrate binding accounts for the ADP-ribosylation defect observed for the ZF PARP7 mutants. Both C243A and C251A PARP7 ZF mutants retained auto-ADP-ribosylation as measured by AF1521 pull-down, indicating they were enzymatically active. Furthermore, the C243A PARP7 mutant co-immunoprecipitated with AR suggesting that the ZF domain is dispensable in the context of full-length PARP7 binding to AR. A follow-up study showed that an intact ZF domain was also dispensable in the context of the NTR PARP7 binding to AR, and GST pull-down showed that the isolated ZF is not sufficient to interact with AR. As shown by the Matthews group [20,22], we confirmed that the ZF PARP7 mutant displays a defect in terms of PARP7 nuclear localization. Although nuclear localization of PARP7 is critical for AR ADP-ribosylation and can be disrupted by mutations within the ZF, we found that rescuing nuclear localization by fusing the SV40 NLS to the C243A PARP7 mutant did not restore ADP-ribosylation of AR.

The data suggest two models for how the ZF may be contributing to AR ADP-ribosylation. One possibility is that the ZF is playing a protein structural role that helps modulate PARP7 ADP-ribosylation activity toward AR. Although the ZF is dispensable for the interaction between PARP7 and AR, the domain could be forming transient contact sites with AR as the receptor undergoes complex conformational rearrangement upon ligand binding, allowing for productive engagement of the AR ADP-ribosylation sites with the PARP7 catalytic domain.

A second model for the role of the ZF in AR ADP-ribosylation relates to the formation of PARP7 foci in the nucleus. We have confirmed the earlier reports about nuclear PARP7 foci formation [20,23], but also found, strikingly, that the C243A and the NLS-C243A PARP7 mutants had virtually no foci formation, indicating that assembly of PARP7 into liquid droplets requires the ZF. The PARP7 ZF is classified as a Cys_3_His_1_-type ZF which is generally associated with RNA binding [27]. The propensity for PARP7 to form liquid droplets in the nuclei could be due to the ZF functioning as an RNA binding module as well as the presence of an N-terminal intrinsically disordered region predicted by IUPred2A (Figure 2A) [28]. Both intrinsically disordered regions and RNA binding domains have been implicated in liquid droplet assembly [29–31]. Based on our results, an intriguing notion is that the PARP7 ZF mediates assembly of PARP7 into nuclear foci which are linked to AR ADP-ribosylation. For example, the foci may represent large macromolecular complexes that PARP7 forms in order to be able to ADP-ribosylate AR.

Overall, our study sheds light on how PARP7-mediated AR ADP-ribosylation is tightly coupled to ligand binding to AR. In the absence of androgen, AR ADP-ribosylation is undetectable, and this strict dependence of AR ADP-ribosylation on androgen is the result of multiple features within the PARP7/AR signaling axis (Figure 6). First, androgen binding to AR induces an agonist conformation that promotes AR import into the nucleus where co-compartmentalization of the PARP7-AR enzyme-substrate pair can occur. A NLS has been characterized in PARP7 [22], and we and others have observed that PARP7 localizes to the nucleus preferentially [9,20,22,32]. Furthermore, we have shown in this study that nuclear localization of PARP7 is required for AR ADP-ribosylation. Given that androgen binding to AR drives its import into the nucleus [33–35] where PARP7 localizes, it is logical that androgen-dependence of AR ADP-ribosylation is in part explained by the increased concentration of AR in the nucleus for PARP7 to act on. Second, *PARP7* is a direct AR target gene that is induced by androgen treatment [7,8]. We have found that almost no PARP7 is detected by immunoblotting in PCa cells prior to androgen treatment, indicating that active AR signaling is required for PARP7 expression [9]. Third, PARP7 protein is stabilized by androgen signaling [9]. The androgen-dependent stabilization is a post-transcriptional mechanism that works in concert with *PARP7* mRNA induction to increase PARP7 protein level in cells. Critically, PARP7 protein stabilization leads to PARP7 accumulation in the nucleus [9], precisely where AR, its substrate, is now concentrated by androgen-induced nuclear import. Fourth, as demonstrated with the experiments using the AR L860F and T877A mutants, the agonist conformation of AR is required for AR ADP-ribosylation by PARP7. These features illustrate the complex interplay between androgen signaling and activity of a specific PARP family member, and help explain how the AR LBD is used as part of a molecular switch that regulates PARP7/AR signaling axis.

**Figure 5.**
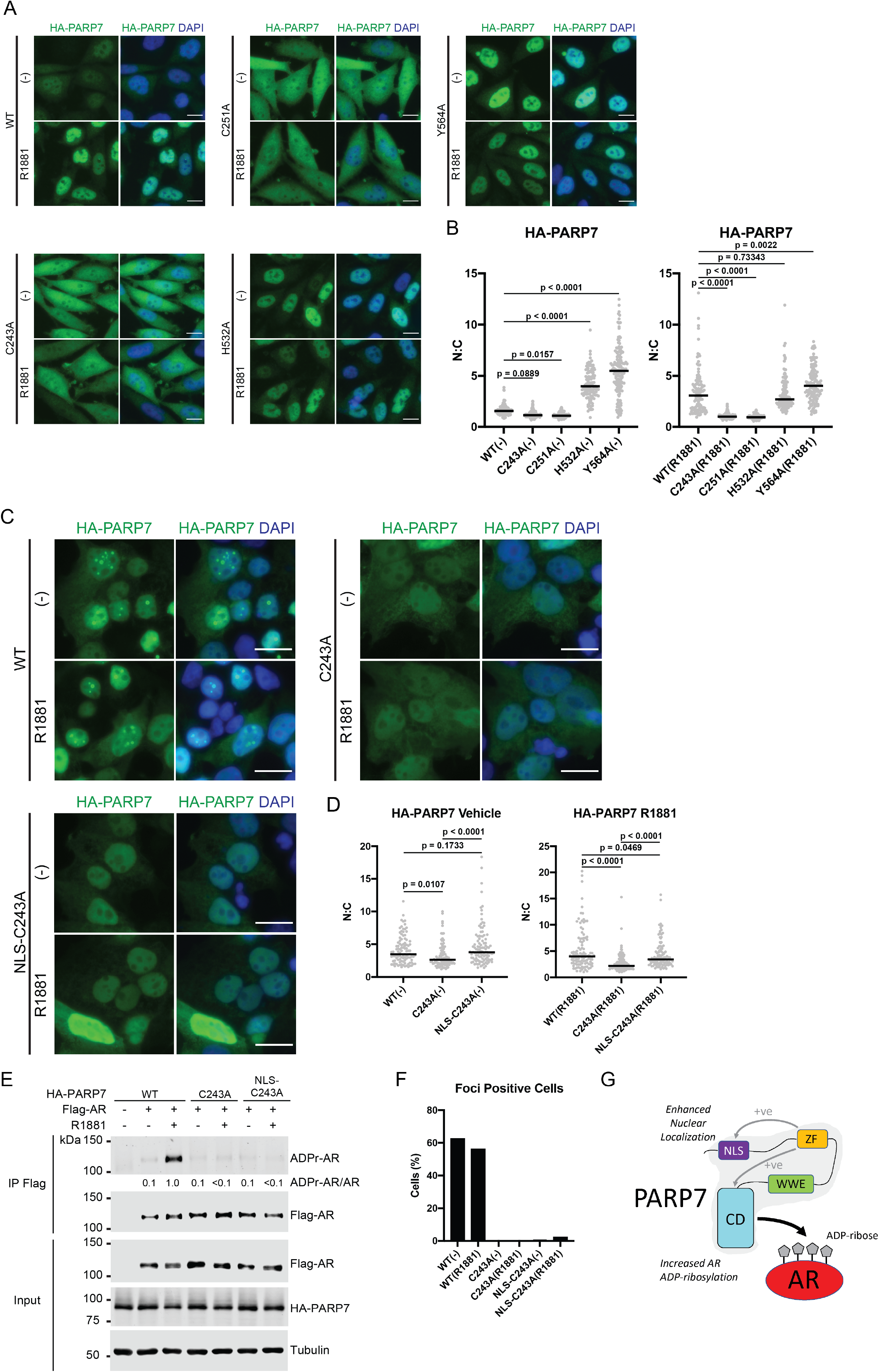
Rescue of the nuclear localization defect for PARP7 zinc finger mutant does not restore AR ADP-ribosylation. (**A**) PC3-Flag-AR cells expressing HA-PARP7 WT or mutant were induced with doxycycline and treated with R1881 similar to Fig 2B. Treated cells were processed for immunofluorescence microscopy. Scale bar = 5 μm. (**B**) Quantification of A). Distribution of HA-PARP7 was quantified as a ratio of nuclear (N) and cytoplasmic (C) signals for vehicle- (left) and R1881-treated (right) groups. At least 100 cells were quantified for each condition, and median N:C ratio (black line) is indicated on the plots. One-way ANOVA with Tukey’s multiple comparison test was conducted to determine statistical significance. (**C**) HEK293T cells were transfected with Flag-AR along with HA-PARP7 WT, C243A, or NLS-C243A (SV40 NLS fused to C243A PARP7 mutant). After 24 h, cells were treated with R1881 for 8 h and processed for immunofluorescence microscopy. Scale bar = 5 μm. (**D**) Quantification of C). Distribution of HA-PARP7 was quantified as a ratio of nuclear (N) and cytoplasmic (C) signals for vehicle (left) and R1881-treated (right) groups similar to B). At least 100 cells were quantified for each condition, and median N:C ratio (black line) is indicated on the plots. One-way ANOVA with Tukey’s multiple comparison test was conducted to determine statistical significance. (**E**) HEK293T cells were transfected with Flag-AR along with HA-PARP7 WT, C243A, or NLS-C243A and treated as in C). Flag-AR was immunoprecipitated from treated cells and analyzed for AR ADP-ribosylation. ADP-ribosylated AR (ADPr-AR) was quantified as a ratio of ADPr-AR to AR. Ratios are displayed relative to the WT + R1881 lane (set to 1.0). (**F**) Quantification of C). HA-PARP7 foci-containing cells were counted (at least 200 cells per condition) and expressed as a percent of total cells counted. (**G**) Working model summarizing the ZF contribution to PARP7 activity. The intact ZF could enahnce steady state nuclear localization by promoting recognition of the NLS by the import machinery, or through a nuclear retention mechanism. The contribution to AR ADP-ribosylation appears to be independent of PARP7 nuclear localization; thus, the PARP7 ZF is envisioned to modulate some aspect of PARP7 structure that affects substrate modification with ADP-ribose. These effects are lost in Cys-to-Ala mutants of PARP7. NLS = nuclear localization signal, ZF = zinc finger, WWE = tryptophan-tryptophan-glutamate, CD = catalytic domain, +ve = positive.

**Figure 6.**
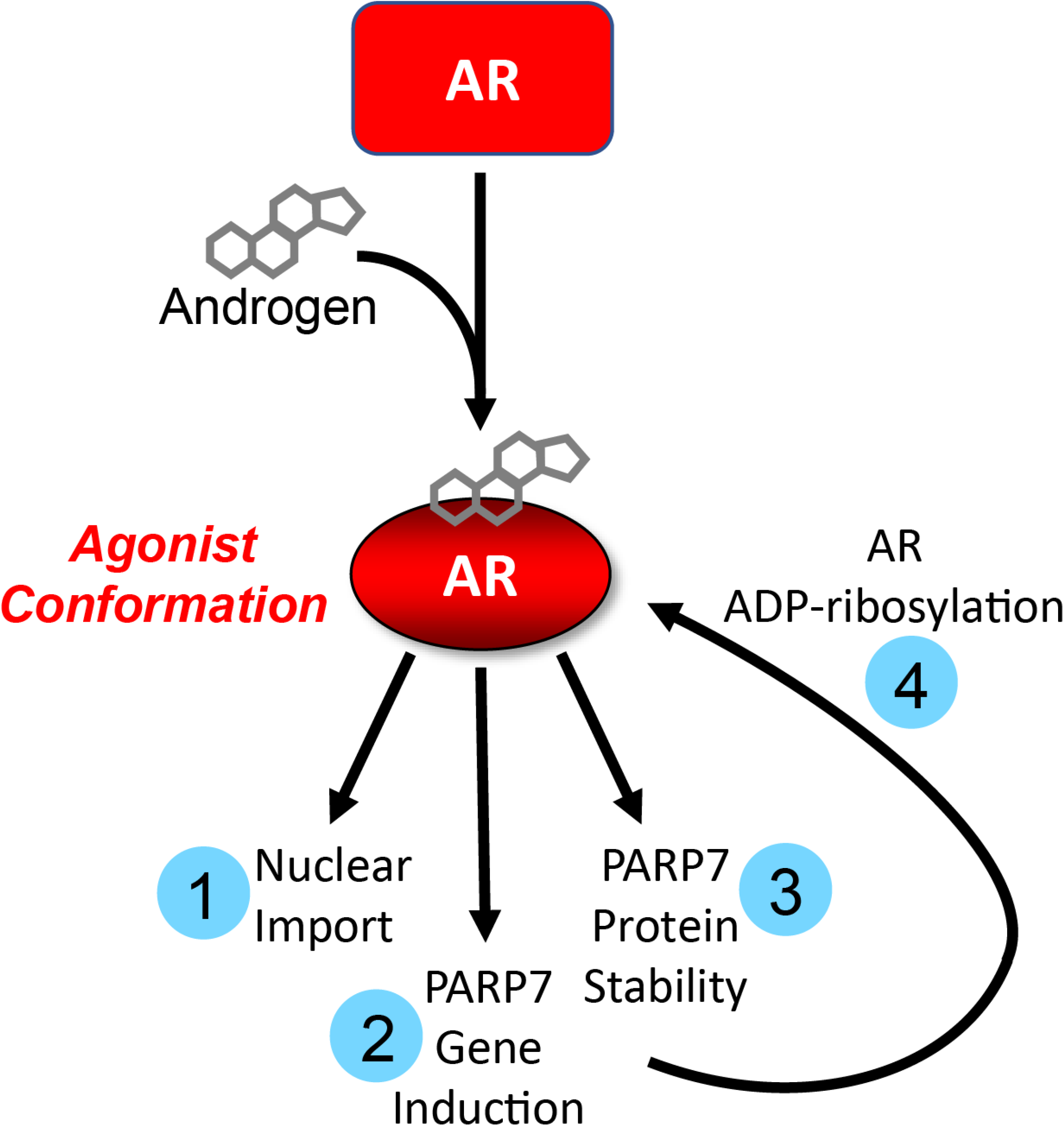
Multiple androgen-dependent mechanisms regulate AR ADP-ribosylation by PARP7. Androgen binding induces an agonist conformation in AR that supports AR ADP-ribosylation through the following: 1) AR import into the nucleus is promoted where PARP7 colocalizes, 2) AR is activated as a transcription factor to induce PARP7 gene expression, 3) AR-dependent transcription leads to PARP7 protein stabilization and accumulation in the nucleus, and 4) PARP7 ADP-ribosylates the agonist conformation of AR.

## Materials and methods

### Plasmid DNA

N-terminally 3xHA-tagged PARP7 WT, N-terminal region (amino acids: 1-456; NTR), and catalytic domain (amino acids: 457-657; CD) were cloned into the pKH3 vector. SDM was conducted on pKH3/HA-PARP7 and pKH3/HA-PARP7 NTR to generate the following expression vectors: pKH3/HA-PARP7 C243A, pKH3/HA-PARP7 H532A, and pKH3/HA-PARP7 NTR-C243A. Additionally, SDM was used to introduce K41A/K42A/K43A substitutions in the TetON-HA-PARP7 lentiviral vector [9] to generate the TetON-HA-PARP7 AAA lentiviral vector. Annealed oligo cloning strategy was used to insert either the SV40 NLS (PKKKRKV) or c-Abl NES (EAINKLESNLRELQICPAT) between the HA tag and the PARP7 CDS to generate the following vectors: TetON-HA-PARP7 NLS, TetON-HA-PARP7 NES, TetON-HA-PARP7 NLS-AAA, TetON-HA-PARP7 NES-AAA, and pKH3/HA-PARP7 NLS-C243A. pCDNA3/Flag-AR was generated previously [36], and SDM was conducted on the vector to introduce the L860F substitution. pcDNA3/Flag-AR T878A was generated previously [33]. PARP7 zinc finger (ZF) domain (amino acids: 232-269) was cloned into pGEX-vector for expression of the GST-fusion protein in *E. coli.* SDM was conducted on pGEX/GST-PARP7-ZF to introduce the C243A substitution in the zinc finger domain (pGEX/GST-PARP7-ZF-C243A)

### Chemical Reagents

The following drugs were used in this study: R1881 (methyltrienolone; used at 2 nM) (PerkinElmer, Inc., Waltham, MA, USA) and hydroxyflutamide (HO-flutamide; used at 1 μM) (Sigma-Aldrich, St. Louis, MO, USA).

### Antibodies

The following primary antibodies were used for this study: anti-AR (custom rabbit polyclonal against AR amino acid: 1-21 or amino acid: 656 to 669; Cocalico Biologicals, Inc., Stevens, PA, USA), anti-HA (mouse monoclonal clone 16B12; Covance, Princeton, NJ, USA), anti-tubulin (mouse monoclonal clone TUB-1A2; Sigma-Aldrich), anti-DTX3L (custom rabbit polyclonal against DTX3L catalytic domain; Cocalico Biologicals, Inc.), and anti-PARP9 (custom rabbit polyclonal against PARP9 catalytic domain; Cocalico Biologicals, Inc.). The following secondary antibodies were used for immunoblotting: IRDye^®^ 800-conjugated goat anti-mouse IgG (610-132-121; Rockland Immunochemicals, Inc., Limerick, PA, USA) and AlexaFluor^®^ 680-conjugated donkey anti-rabbit IgG (A10043; Thermo Fisher Scientific, Waltham, MA, USA). For immunofluorescence microscopy, the following secondary antibodies were purchased from Jackson ImmunoResearch Laboratories, Inc. (West Grove, PA, USA): Cy3-conjugated donkey anti-mouse (715-165-151), Cy5-conjugated donkey anti-rabbit (711-175-152), and AlexaFluor^®^ 488-conjugated donkey anti-mouse (715-545-150) antibody.

### Cell culture and transfections

PC3-Flag-AR and PC3-Flag-AR/TetON-HA-PARP7 (WT, C243A, C251A, H532A, and Y564A) cell lines were generated previously [9,37]. PC3-Flag-AR/TetON-HA-PARP7 NLS, PC3-Flag-AR/TetON-HA-PARP7 NES, PC3-Flag-AR/TetON-HA-PARP7 AAA, PC3-Flag-AR/TetON-HA-PARP7 NLS-AAA, and PC3-Flag-AR/TetON-HA-PARP7 NES-AAA cell lines were derived from PC3-Flag-AR via lentivirus transduction. All PC3 derived cells were grown in RPMI 1640 medium supplemented with 5% fetal bovine serum (SH30396.03HI; Cytiva, Marlborough, MA, USA) and 100 U/mL penicillin/streptomycin (Thermo Fisher Scientific). Puromycin (1 μg/mL) was added to culture medium to maintain selection for cell lines carrying the TetON vector. HEK293T cells were grown in DMEM/F12 (1:1) medium supplemented with 5% fetal bovine serum (Cytiva) and 100 U/mL penicillin/streptomycin (Thermo Fisher Scientific). All cells were grown at 37°C with 5% CO_2_. Transient transfections into HEK293T cells were conducted using Lipofectamine 3000 (Thermo Fisher Scientific) according to manufacturer’s protocol.

### Detection of ADP-ribosylation

AF1521 is a macrodomain protein from the bacteria *Archaeoglobus fulgidus* that specifically binds to ADP-ribose [38] and was utilized in pull-down or as a direct blotting reagent for detection of ADP-ribosylation as described in [39]. Treated cells were lysed in 50 mM Tris-HCl pH 7.5, 150 mM NaCl, 0.5% Triton X-100 (v/v), 1 μg/mL aprotinin, 1 μg/mL leupeptin, 1 μg/mL pepstatin, 1 mM PMSF, and 1 mM DTT and incubated with magnetic glutathione beads (L00327; GenScript Biotech, Piscataway, NJ, USA) loaded with recombinant GST-tagged AF1521 for 2 to 4 hours at 4°C with rotation. Beads were washed five times with wash buffer (50 mM Tris-HCl pH 7.5, 150 mM NaCl, 0.1% Triton X-100 (v/v), and 1 mM DTT) and resuspended in 1× sample buffer for subsequent analysis by SDS-PAGE and immunoblotting. For direct blotting, recombinant GST-tagged tandem AF1521 was fluorescently labeled using IRDye^®^ 800CW Protein Labeling Kit (LI-COR Biosciences, Lincoln, NE, USA). Prepared samples were run on SDS-PAGE, and proteins were immobilized on nitrocellulose membranes. After blocking for at least 1 h in milk solution (5% nonfat dry milk (w/v)/1× PBS with 0.15% Tween 20 (v/v) (1× PBST)), membranes were probed with the fluorescently-labeled AF1521 to detect ADP-ribosylated AR, and images were acquired using an Odyssey^®^ CLx system (LI-COR Biosciences).

### Co-immunoprecipitation

For co-immunoprecipitations from treated PC3-Flag-AR/TetON-HA-PARP7 cells, collected cell pellets were lysed in 50 mM Tris-HCl pH 7.5, 150 mM NaCl, 0.5% Triton X-100 (v/v), 1 μg/mL aprotinin, 1 μg/mL leupeptin, 1 μg/mL pepstatin, 1 mM PMSF, and 1 mM DTT. For co-immunoprecipitations from treated HEK293T cells, collected cell pellets were lysed in 20 mM HEPES pH 7.5, 200 mM NaCl, 0.5% NP-40, 1 μg/mL aprotinin, 1 μg/mL leupeptin, 1 μg/mL pepstatin, 1 mM PMSF, and 1 mM DTT. Clarified cell extract were applied to magnetic anti-Flag M2 beads (M8823; Sigma-Aldrich) and incubated for 2 hours at 4°C with rotation. Beads were washed five times with the following wash buffers: 50 mM Tris-HCl pH 7.5, 150 mM NaCl, 0.1% Triton X-100 (v/v), and 1 mM DTT for co-immunoprecipitations from PC3-Flag-AR/TetON-HA-PARP7 cells, and 20 mM HEPES pH 7.5, 200 mM NaCl, 0.1% NP-40, and 1 mM DTT for co-immunoprecipitations from HEK293T cells. Washed beads were resuspended in 1× sample buffer and analyzed by SDS-PAGE and immunoblotting.

### GST-PARP7-zinc finger pull-down

GST-tagged PARP7 zinc finger WT and C243A proteins were recombinantly expressed in *E. coli* and purified as described in [39] with the following modifications: 1) induction of GST-tagged proteins in *E. coli* was done in the presence of 50 μM ZnCl_2_ in the culture medium, 2) lysis and elution buffers were supplemented with 1 μM ZnCl_2_, and 3) after purification, GST-tagged proteins were dialyzed in 1× PBS, 1 μM ZnCl_2_, and 2 mM DTT. Magnetic glutathione beads (L00327; GenScript Biotech) were loaded with the purified PARP7 zinc finger proteins, and subsequently incubated with extracts prepared from cell pellets lysed in 20 mM HEPES pH 7.5, 200 mM NaCl, 0.5% NP-40, 1 μg/mL aprotinin, 1 μg/mL leupeptin, 1 μg/mL pepstatin, 1 mM PMSF, 1 mM DTT, 100 U/mL RNase inhibitor, and 2 nM R1881. After 2 h incubation at 4°C with rotation, beads were washed four times with wash buffer (20 mM HEPES pH 7.5, 200 mM NaCl, 0.5% NP-40, 1 mM DTT, 100 U/mL RNase inhibitor, and 2 nM R1881) and resuspended in 1× sample buffer for analysis by SDS-PAGE and immunoblotting.

### SDS-PAGE and immunoblotting

Samples were prepared in 1× sample buffer and analyzed by SDS-PAGE. After transfer, nitrocellulose membranes were blocked for 1 h in blocking solution (5% nonfat dry milk (w/v)/1× PBS with 0.15% Tween 20 (v/v) (1× PBST)), followed by primary and secondary antibody incubations. Membranes were washed between each step with 1× PBST. Fluorescent signal was detected using an Odyssey^®^ CLx imaging system (LI-COR Biosciences), and quantification was done using the Image Studio Lite version 5.2.5 (LI-COR Biosciences).

### Immunofluorescence microscopy

Coverslips were processed for immunofluorescence microscopy as described in [9]. Cells were seeded onto glass coverslips at least 48 h before processing. Treated cells were fixed with 3.75% formaldehyde/1× PBS for 15 minutes, permeabilized with 0.2% Triton X-100/1 × PBS for five minutes, and blocked for 1 h at room temperature in blocking buffer (2% BSA (w/v)/1× PBS). Coverslips were incubated in primary antibody overnight at 4°C, and after washes, incubated in secondary antibody for 1 h at room temperature. Before mounting, washed coverslips were briefly incubated in DAPI to stain for nuclei, followed by a rinsing step with deioinized water. Coverslips were mounted on glass slides with VectaShield (Vector Laboratories, Burlingame, CA, USA). Images were acquired using a Nikon Eclipse Ni-U microscope (Nikon Instruments, Inc., Melville, NY, USA) equipped with a DS-Qi1Mc camera at 40× objective. All images were processed using Adobe Photoshop version 21.2.2 (Adobe Inc., San Jose, CA, USA) and Fiji ImageJ version 2.0.0. HA-PARP7 cellular distribution was quantified as a ratio of nuclear (N) to cytoplasmic (C) signal as described previously [40]. N:C ratio was calculated from the background-corrected mean signal intensities of regions of interest outlined in the nucleus and the cytoplasm. At least 100 cells were quantified for each condition.

### RT-qPCR

To isolate RNA, treated cells were processed using the RNeasy kit (QIAGEN, Hilden, Germany). cDNA was synthesized from 1 μg of isolated RNA using iScript cDNA synthesis kit (Bio-Rad Laboratories, Hercules, CA, USA). RT-qPCR was carried out using SensiMix™ SYBR^®^ and Fluorescein kit (QT615-05; Bioline, London, United Kingdom). The following primers were used: *MYBPC1* (5’-GTCGCTCTCACATGGACTCC-3’ and 5’-AATGGTGGCACTGGTTCGAT-3’) and *GUS* (5’-CCGACTTCTCTGACAACCGACG-3’ and 5’-AGCCGACAAAATGCCGCAGACG-3’). Gene expression was normalized against the housekeeping gene *GUS*, and the mean and standard deviation were calculated from three biological replicates.

### Statistical Analysis

All statistical analysis was conducted in GraphPad Prism version 9.0.1 software (GraphPad Software, San Diego, CA, USA). Statistical significance was determined using a one-way ANOVA with Tukey’s multiple comparison test as appropriate.

## Data Availability Statement

Source data available upon request.

